# Effect of host genetics on the gut microbiome in 7,738 participants of the Dutch Microbiome Project

**DOI:** 10.1101/2020.12.09.417642

**Authors:** E.A. Lopera-Maya, A. Kurilshikov, A. van der Graaf, S. Hu, S. Andreu-Sánchez, L. Chen, A. Vich Vila, R. Gacesa, T. Sinha, V. Collij, M.A.Y. Klaassen, L.A. Bolte, M.F. Brandao Gois, P.B.T. Neerincx, M.A. Swertz, LifeLines Cohort Study, H.J.M. Harmsen, C. Wijmenga, J. Fu, R.K. Weersma, A. Zhernakova, S. Sanna

## Abstract

Host genetics are known to influence the gut microbiome, yet their role remains poorly understood. To robustly characterize these effects, we performed a genome-wide association study of 207 taxa and 205 pathways representing microbial composition and function within the Dutch Microbiome Project, a population cohort of 7,738 individuals from the northern Netherlands. Two robust, study-wide significant (*p*<1.89×10^-10^) signals near the *LCT* and *ABO* genes were found to affect multiple microbial taxa and pathways, and were replicated in two independent cohorts. The *LCT* locus associations were modulated by lactose intake, while those at *ABO* reflected participant secretor status determined by *FUT2* genotype. Eighteen other loci showed suggestive evidence (*p*<5×10^-8^) of association with microbial taxa and pathways. At a more lenient threshold, the number of loci identified strongly correlated with trait heritability, suggesting that much larger sample sizes are needed to elucidate the remaining effects of host genetics on the gut microbiome.

## Introduction

The human intestinal microbial community contains trillions of microorganisms that play an important role in maintaining normal gut function and immune homeostasis^1^ . Emerging evidence has shown that alterations in gut microbial composition are associated with the pathogenesis of many human diseases, including gastrointestinal disorders, metabolic syndrome, cardiovascular diseases and other conditions^2, 3^ .

The gut microbiome is influenced by many factors, such as environmental factors including diet and medication usag ^4, 5^, but also by host genetics. Heritability studies in twins and families have highlighted that human genetics contributes to gut microbial variation, showing heritability estimates ranging from 1.9% to 8.1%^6, 7^ . This observation encouraged the first efforts to identify genomic loci that influence gut microbiota through genome-wide association studies (GWAS). These early gut microbiome GWAS identified several microbial quantitative trait loci (mbQTL) located in genes related to intestinal mucosal barrier, immune response and drug and food metabolism^8–11^ . However, reproducibility of these findings has been limited by differences in data processing methodologies, modest sample sizes and strong environmental effects, which, taken together, limit the detection of robust host genetic associations^12^ . A recent large-scale genome-wide meta-analysis of 24 cohorts replicated the association between *Bifidobacterium* abundance and the lactase (*LCT*) gene locus^13^, which had previously only been reported in single-cohort studies^6, 14^ . Other suggestive mbQTLs identified in this broad meta-analysis were proportional to heritability estimates from independent twin studies, indicating that additional loci found at lenient levels of significance are likely to be real but that larger sample sizes are needed to reach sufficient statistical power^13^. Nonetheless, meta-analyses of mbQTL studies still lack resolution due to the high levels of heterogeneity between cohorts, which can reduce power in meta-analysis. On top of this, many existing cohorts rely on 16S rRNA measurements, but this method does not allow for identification at bacterial species-level resolution or of bacterial pathway abundances. Indeed, measuring both species- and pathway-level abundances is essential for a further understanding of the function of an individual’s microbiome because two bacterial species can have very different functions in the gut microbiome, yet pathways may be shared across distant microbial species and have roughly the same biological effect^15^ . mbQTL studies using shotgun metagenomic sequencing in large population cohorts are therefore needed to overcome the high variability in microbiome definition and reveal more robust and refined associations.

To enable a broader and deeper understanding of host‒microbiota interactions, we used shotgun metagenomic sequencing on feces from 7,738 individuals of the Dutch Microbiome Project (DMP)^16^ and matched their imputed genotypes to differences in taxa and pathway abundances. By comparing our results with summary statistics from other independent studies^13, 17, 18^, we reveal novel host‒microbiota interactions. Furthermore, we explore the impact of both intrinsic and environmental factors in confounding or modulating these genetic effects and identify diet-dependent host‒microbiota interactions. We further assess the potential causal relationships between the gut microbiome and dietary habits, biomarkers and disease, using Mendelian randomization. Finally, we carry out a power analysis that shows how microbiome studies, even at the current sample size, are not well powered to reveal the complex genetic architecture by which host genetics can regulate the gut microbiome.

## Results

### Genome-wide association identifies known and novel associations with several bacterial taxonomies and bacterial pathways

We investigated 5.5 million common (minor allele frequency (MAF) > 0.05) genetic variants on all autosomes and the X chromosome, using linear mixed models^19^ to test for additive genetic effects on 207 taxa and 205 bacterial pathways in 7,738 individuals from the DMP cohort (**Methods**)^19^. Our quality control steps did not detect evidence for test statistic inflation (median genomic lambda 1.003 for taxa and 1.01 for pathways). We identified 33 independent SNP‒trait associations at 20 loci at a genome-wide *p-*value threshold of 5×10^-8^ (**Figure 1, Supplementary Table 1**). Two loci passed the more stringent study-wide threshold of 1.89×10^-10^ that accounts for the number of independent tests performed (**Methods**). The other 18 loci were associated at *p-* values between 5×10^-8^ and 1.89×10^-10^, with 4 associated to taxa and 14 to pathways. We refer to these 18 loci as “suggestive” and discuss them below.

**Figure 1.**
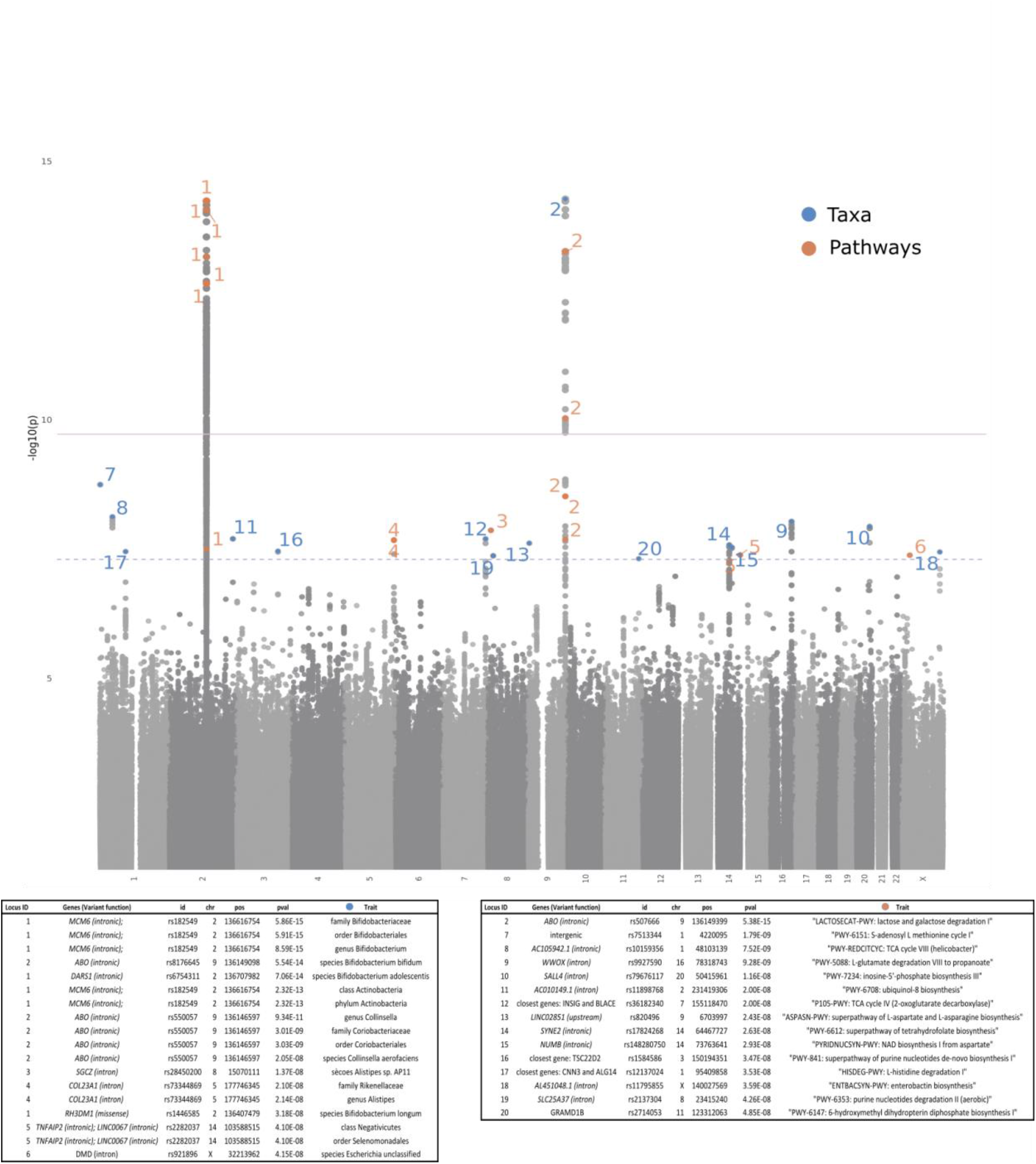
Genome-wide association scan results. Manhattan plot of host genomic associations to bacterial taxa and bacterial pathway abundances with at least one genome-wide significant association (*p* < 5x10^-8^). Y-axis shows the -log10 transformation of the association *p-*value observed at each tested variant. X-axis shows the genomic position of variants. The thresholds of study-wide (*p* = 1.89×10^-10^) and genome-wide (*p* = 5x10^-8^) significance are shown as horizontal red and blue lines, respectively. Independent SNP‒ trait associations reaching genome-wide significance are listed in the tables and labelled on the Manhattan plot. The colors of associated hits indicate whether they represent an association with taxonomy (orange) or a bacterial pathway (blue).

The strongest signal was seen at rs182549 located in an intron of *MCM6*, a perfect proxy of rs4988235 (r^2^ = 1, 1000Genome European populations), a variant known to regulate the *LCT* gene and responsible for lactase persistence in adults (ClinVar accession RCV000008124). The T allele of rs182549, which confers lactase persistence through a dominant model of inheritance, was found to decrease the abundance of species *Bifidobacterium adolescentis* (*p* = 7.6×10^-14^) and *Bifidobacterium longum (p* = 3.2x10^-08^) as well as those of higher-level taxa (phyla *Actinobacteria p*= 2.3×10^-13^; class Actinobacteria, *p* = 2.3×10^-13^; order Bifidobacteriales*, p* = 5.9×10^-15^; family Bifidobacteriaceae, *p* = 5.86×10^-15^; genus *Bifidobacterium, p* = 8.59×10^-15^). Associations at this locus were also seen for other taxa of the same genus, albeit at lower levels of significance (*Bifidobacterium catenulatum*, *p* = 3.9×10^-5^), and for species of the *Collinsella* genus (**Figure 2A**). The association with *Bifidobacterium* at the *LCT* locus has been previously described in Dutch, UK and US cohorts^6, 8, 14^, as well as in a recent large-scale meta-analysis^13^. The presence of the *LCT* association across several taxa indicates that this locus has a wide-ranging effect on microbiome composition.

**Figure 2.**
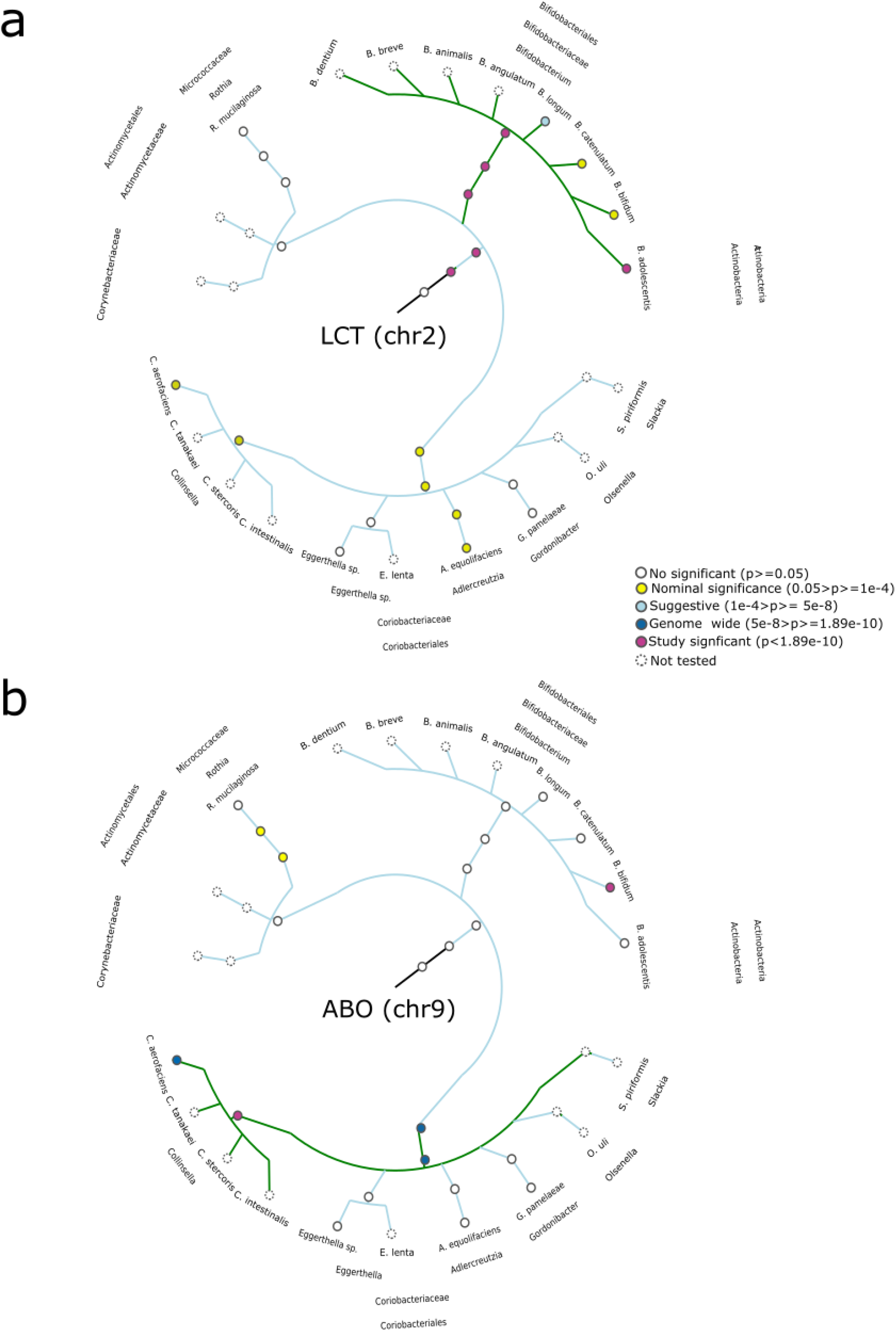
Phylogenetic tree of taxonomic relations between bacteria of the class Actinobacteria and their associations with host genetics. Each node shows a taxonomic level (from outside to inside: phylogenetic group, phylum, class, order, family, genus and species). Note that branch lengths do not represent phylogenetic distance. Inner labels represent genetic locus. External labels represent the clade. Nodes with dotted line type indicate that the GWAS for that taxa was not performed . Node color corresponds to different levels of significance as described in the legend. **a)** depicts associations detected at the *MCM6*/*LCT* locus, using as the most significant *p*-value observed for rs4988235 and rs182549. **b)** depicts associations to the *ABO* locus, using the most significant *p*-value observed for rs8176645 and rs550057.

The second locus that passed study-wide significance consists of genetic variants near the *ABO* gene. *ABO* encodes the BGAT protein, a histo-blood group ABO system transferase. Associations found at this locus include species *B. bifidum* (rs8176645, *p* = 5.5×10^-15^) and *Collinsella aerofaciens* (rs550057, *p* = 2.0×10^-8^, *r*^2^ = 0.59 with rs8176645 in 1000 Genomes Europeans) and higher order taxa (rs550057, genera *Collinsella, p* = 9.3×10^-11^; family Coriobacteriaceae, *p* = 3.01×10^-9^; order Coriobacteriales, *p* = 3.03×10^-9^) (**Figure 2B**). Interestingly, the metabolic pathway representing the bacterial degradation of lactose and galactose is also associated to the *ABO* locus (Metacyc ID: “*LACTOSECAT-PWY: lactose and galactose degradation I*”, rs507666, *p* = 5.38×10^-15^). The association at this locus with the genus *Collinsella* and with metabolic pathway *LACTOSECAT-PWY* have been recently described^18, 20^. To our knowledge, the associations with *B. bifidum* and *C. aerofaciens* are novel.

### Association at *LCT* reflects the lactase persistence inheritance model and affects multiple taxonomic levels and bacterial pathways

Given that lactose tolerance is inherited in a dominant fashion, we tested the associations found in this locus using a dominant model for SNP rs182549 and thereby compared lactase persistent (LP) to lactose intolerant (LI) individuals. Indeed, all seven taxa associated to the *LCT* locus at genome-wide significance showed a stronger association signal when we used a dominant model for the alternative allele (all associations *p* < 2×10^-27^), with decreased taxa abundance in LP individuals (**Supplementary Table 2**). We further tested the other 200 taxa for this SNP using a dominant transmission model. Intriguingly, we also observed suggestive association (*p* < 1×10^-4^) with rs182549 for taxa that were associated to *ABO* locus in our GWAS analysis (genera *Collinsella* and species *B. bifidum* and *C. aerofaciens)* and for the species *Roseburia inulinivorans* of the family Lachnospiraceae (**Supplementary Table 2**). With the exception of *Roseburia inulinivorans,* there was a consistent direction of effect across the associated taxa, with increased abundance in LI compared to LP individuals (**Figure 3**).

**Figure 3.**
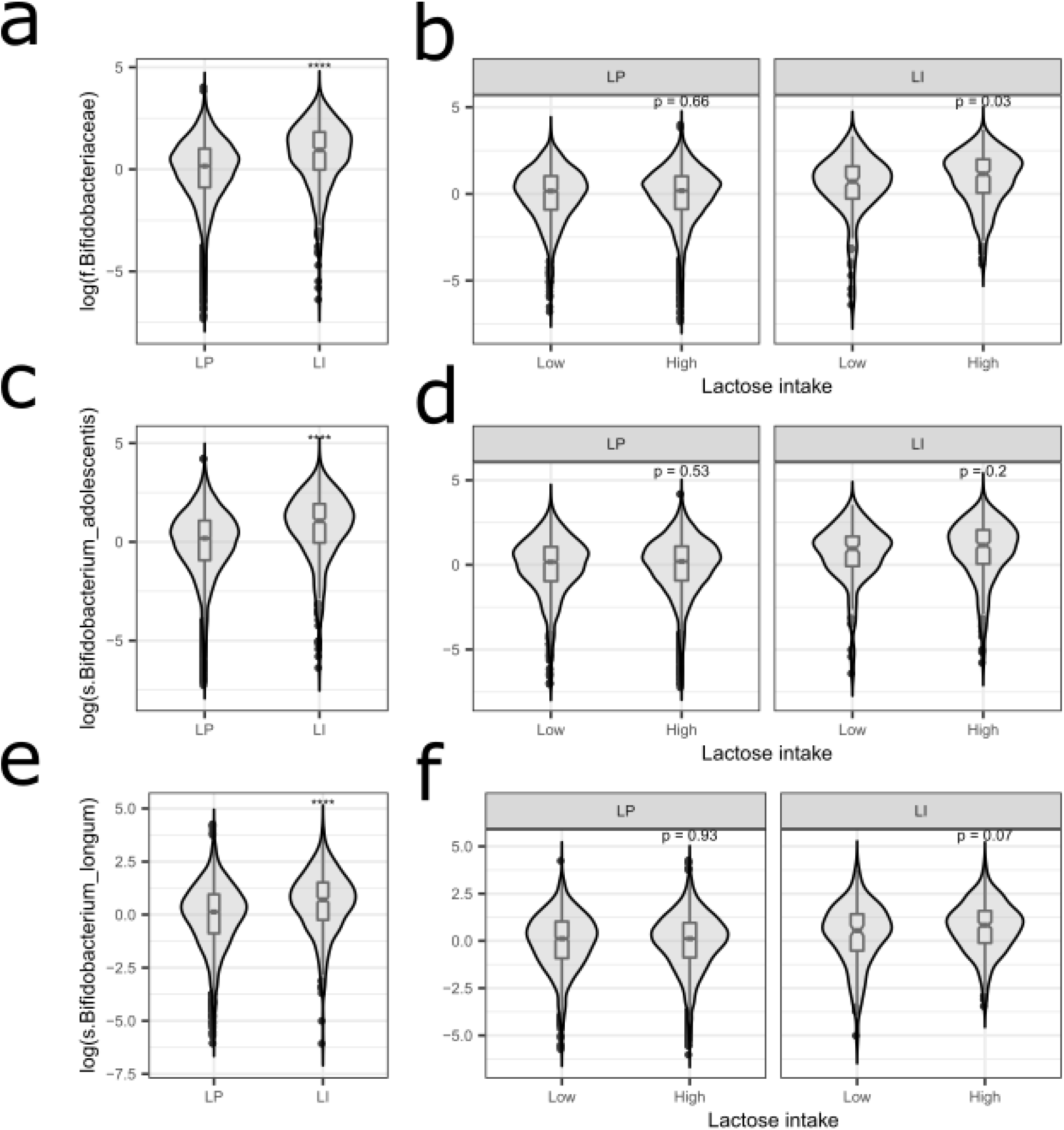
Association at the *LCT* locus and interaction with lactose intake. Relative abundance of taxa found associated with SNPs at the *LCT* locus compared between Lactase Persistent (LP, rs182549 C/T or T/T) and Lactose Intolerant (LI, rs182549 C/C) participants and among individuals assuming low or high daily lactose intake levels (lactose intake was corrected for daily calorie consumption). Relative abundance is displayed with a violin plot, where each inner box represents the first and third quartiles of the distribution and the middle line represents the median. Left panels are different relative abundances for different taxa: **a**) family Bifidobacteriaceae in LP (n = 6,809) and LI (n = 443), **c**) species *Bifidobacterium adolescentis* in LP (n = 5928) and LI (n = 400), **e**) species *Bifidobacterium longum* in LP (n = 6239) and LI (n = 411). Right panels are comparisons of abundance between lactose intake levels, low (<first quartile) and high (≥ first quartile), stratified by lactose persistence: **b**) Bifidobactericeae in 5,081 LP samples with low (n = 1374) or high (n = 4427) lactose intake and in 376 LI samples with low (n = 130) or high (n = 246) lactose intake, **d**) species *Bifidobacterium adolescentis* in 5,017 LP samples with low (n = 1204) or high (n = 3813) lactose intake and in 339 LI samples with low (n = 118) or high (n = 221) lactose intake and **f**) species *Bifidobacterium longum* in 5,301 LP samples with low (n = 1,269) or high (n = 4,032) lactose intake and in 345 LI samples with low (n = 114) or high (n = 231) lactose intake. *P*-values are two-sided *p*-values obtained with a Wilcoxon rank test. **** *p* ≤ 0.0001.

Furthermore, when analyzing LI and LP individuals separately in the bacterial pathways, rs182549 was also associated with an increased abundance of the “*LACTOSECAT-PWY”* bacterial pathway in LI individuals (effect = +0.300 in standard deviation (SD) units, standard error (SE) = 0.49, *p* = 1.02×10^-9^). This is not surprising given that the taxa in our dataset that were most highly correlated with this pathway are of the class Actinobacteria and species *B. adolescentis (*Spearman correlation of 0.73 and 0.69, respectively), which are both associated with SNPs near *LCT*.

### Associations at *ABO* are dependent on secretor status

To further understand the mechanisms underlying the association signal at the *ABO* locus, we derived blood group types based on the genotype status of three genetic variants (see **Methods**). The majority of the individuals were either type A (40%) or type O (48%), reflecting estimates from previous studies ^21^. We observed that the associations at this locus were driven by individuals producing either A or B antigens compared to individuals with blood type O. The abundances of the genera *Collinsella* and the metabolic pathway “*LACTOSECAT PWY*” were higher in individuals producing A or AB antigens (Wilcoxon test for *Collinsella*, blood type O vs. A: *p* = 2.8×10^-9^, O vs. AB: *p* = 0.11; Wilcoxon test for metabolic pathway, O vs. A: *p* = 5×10^-14^, O vs. AB: *p* = 0.038). In contrast, the abundance of *B. bifidum* was lower compared to individuals with blood type O (Wilcoxon test blood type, A vs. O: *p* = 2.3×10^-14^, B vs. O: *p* = 0.007, AB vs. O: *p* = 0.006) (**Figure 4**). Secretion of the antigens produced by the A or B blood groups in body fluids, including the gut mucosa, is dependent on the individual’s secretor status. Secretor status is controlled by the fucosyltransferase 2 (*FUT2*) gene in a recessive manner and determined by nonsense mutation rs601338 (OMIM 182100)^22^. Non-secretors carry the A/A genotype, which leads to a non-functional *FUT2* gene and inability to present ABO antigens on the intestinal mucosa. We observed that the associations of taxa and bacterial pathways with blood types were only present in secretors and absent in non-secretors (**Figure 4**). This indicates that the association at the *ABO* locus is driven by the exposure of blood antigens to gut bacteria. These results recapitulate recent evidence for an mbQTL effect of this locus on the *Collinsella* genus and on other bacteria reported for a Finnish population cohort ^18^. Associations of a functional variant in the *FUT2* gene to other bacterial taxa were observed in a recent meta-analysis, but none of these bacterial taxa or pathways showed significant association for this locus in our cohort ^13^.

**Figure 4.**
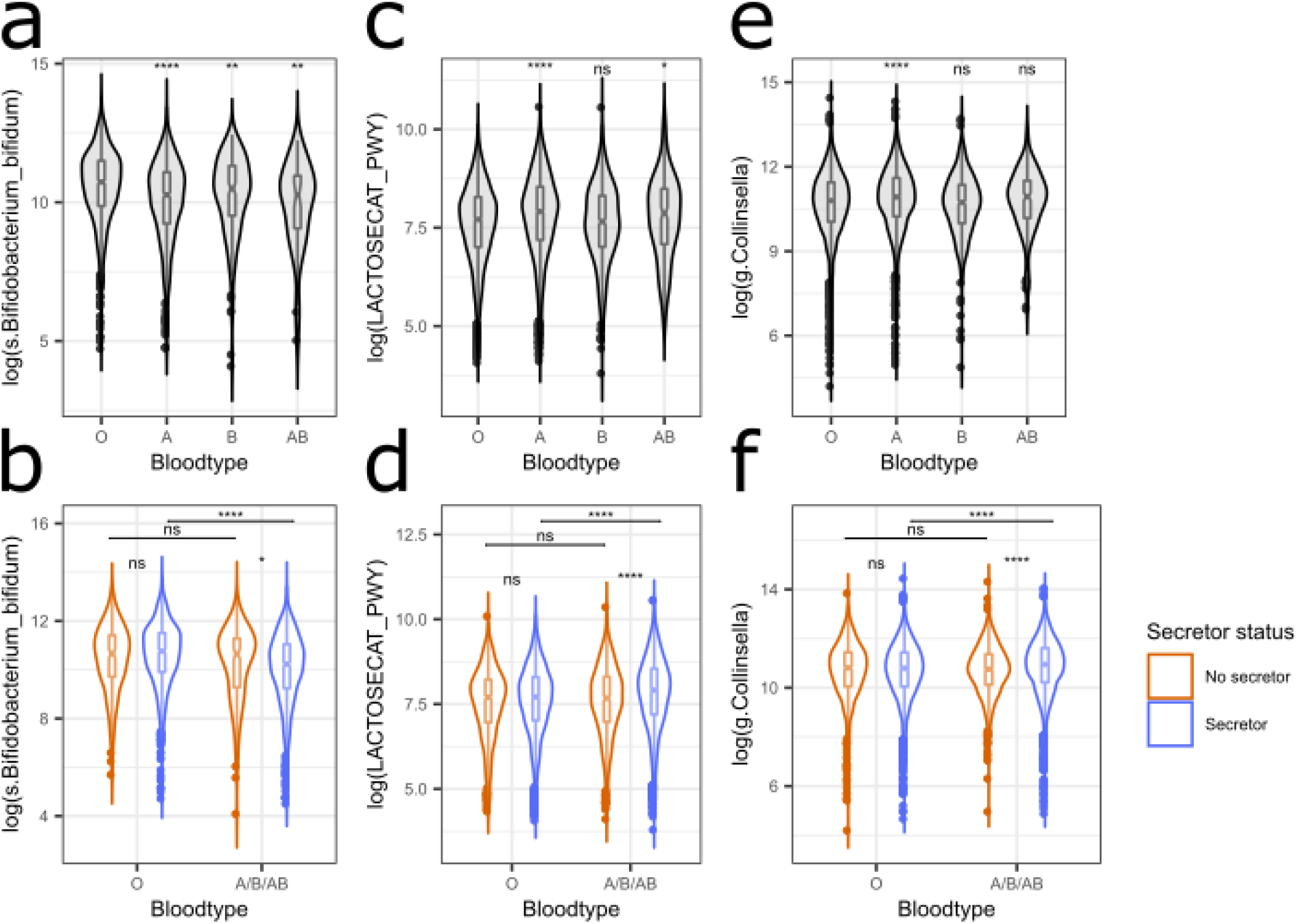
Association with blood types and interaction with *FUT2*. Relative abundance of features found to be associated with SNPs at the *ABO* locus are compared across the inferred blood types (O, A, B and AB) with O as reference group (panels (**a**), (**b**) and (**c**)), and between secretors (blue) and non-secretors (orange) of blood antigens, stratified again by the blood types with presence (A/B/AB) or absence (O) of terminal sugars (panels (**d**), (**e**) and (**f**)). Relative abundance is shown using a violin plot in which the inner box represents the first and third quartiles of the distribution and the middle line represents the median. **a**) species *Bifidobacterium Bifidum* in blood types O (n = 1,058), A (n = 757), B (n = 179) and AB (n = 47) (ANOVA *p* = 6.7×10^-14^). **c**) *LACTOSECAT-PWY* in blood types O (n = 3,470), A (n = 2,923), B (n = 621) and AB (n = 210) (ANOVA *p* = 1.4×10^-13^). **e**) genus *Collinsella* in blood types O (n = 3,462), A (n = 2,906), B (n = 632) and AB (n = 210) (ANOVA *p* = 5.4x10^-9^). **b**) species *Bifidobacterium Bifidum* in blood O antigen secretors (n = 884) and non-secretors (n = 174) and in blood A, B or AB antigen secretors (n = 798) and non-secretors (n = 185). **d**) *LACTOSECAT-PWY* in blood O antigen secretors (n = 2,669) and non-secretors (n = 801) and in blood A, B or AB antigen secretors (n = 2,873) and non-secretors (n = 881). **f**) genus *Collinsella* in blood O antigen secretors (n = 2,664) and non-secretors (n = 798) and in blood A, B or AB antigen secretors (n = 2,864) and non-secretors (n = 884). P-values are two-sided *p*-values obtained from the Wilcoxon ranked test. ns *p* > 0.05, * *p* ≤ 0.05, ** *p* ≤ 0.01, *** *p* ≤ 0.001, **** *p* ≤ 0.0001.

### Genetic associations are not confounded by intrinsic factors but may be modulated by diet

Gut microbiome composition and function is known to be affected by intrinsic factors such as sex or body-mass index (BMI) and by external factors such as diet and medication usage ^4^. We first explored the impact of BMI, medication usage and stool frequency and consistency. None of our genome-wide significant associations were attenuated when including these factors as covariates (**Methods**, all comparisons, Cochran’s Q for difference in effect size *p* > 0.05), indicating that the genetic associations are independent and not confounded by these factors (**Supplementary Table 3a**). Finally, we did not find sex-specific associations for the 33 SNP‒trait associations, although five associations did exhibit a smaller genetic effect in males compared to females (**Supplementary Table 3b**).

We further investigated the effect of diet at the *LCT* and *ABO* loci, taking into account the dominant inheritance model at *LCT* and the dependence on secretor status at *ABO*. From 21 dietary factors derived from a food questionnaire, we considered those previously found to be associated (FDR < 0.05) with the microbial taxa and pathway abundances showing genome-wide association signals in these two loci (see **Methods**)^16^. For the genus *Collinsella* and pathway *LACTOSECAT-PWY*, no dietary factors were found at this FDR threshold, we therefore considered the same diet factors associated to *B. bifidum* in the analyses, given that they showed similar patterns of genetic association. In an analysis that included age, gender, genetic and dietary factors, the dietary factors did not alter the effect of the genetic components, indicating that diet is not a source of bias in these genetic associations (**Supplementary Table 4**). Diet also remained an important factor after including genetics, with four taxa and one pathway associated at *LCT* and *ABO* being affected by at least one dietary factor (*p* < 0.05)(**Supplementary Table 4**), with the maximum of 16 factors found for *B. longum*. Among all the diet factors that showed association with the microbiome features after accounting for the genetic signals (44 diet‒microbiome pairs), we found evidence for a gene‒diet interaction for one bacteria at the *LCT* locus: an increased abundance of Bifidobacteriaceae family in LI individuals who consumed larger amounts of lactose or dairy (interaction term *p*=0.03) (**Figure 4, Supplementary Table 5**), a finding consistent with previous reports ^8, 18^. In contrast, there was no evidence for interaction with diet at the *ABO* locus for either the *B. bifidum* species, the *Collinsella* genus or the bacterial pathway “*LACTOSECAT-PWY*” (interaction term *p* > 0.05) (**Supplementary Table 5)**.

### Association signals reveal a polygenic architecture of microbiome taxa and pathways and replicate in other independent cohorts

Apart from the study-wide significant associations at the *LCT* and *ABO* gene regions, 18 other loci showed suggestive association at *p* < 5×10^-8^ (**Supplementary Table 1**), and none of these have been reported previously. The majority (14) were associated with bacterial pathways, which could not have been directly quantified in studies using 16S rRNA data, the technology predominantly used in microbiome genetic studies to date. These associations included several genomic loci located in genes associated with human immune or metabolic disorders and mainly affected microbial amino acids (3), vitamin (4), purine (3), TCA cycle (2) and enterobactin toxin (1) pathways. These SNP‒pathway associations provide insights into the relationships between host genetics and gut microbial metabolic functions. For instance, SNP rs9927590 in 16q23.1 (in gene *WWOX*), which has been reported to be involved in obesity and stenosis in Crohn’s disease^23, 24^, was associated with a metabolic pathway involved in short chain fatty acid (SCFA) production (*PWY-5088*, *p* = 9.2×10^-9^). SCFAs have been shown to boost intestinal barrier functions and suppress metabolic diseases, including type 1 and type 2 diabetes^17, 25^. A variant (rs2714053) in 11q24.1 (in gene *GRAMD1B*) resulted in lower abundance of bacterial genes involved in the folic acid biosynthesis (*PWY-6147*, *p* = 4.58×10^-8^), and this gene was previously identified to be related to ulcerative colitis in a Polish cohort^26^. Another example is the association nearf the type 2 diabetes‒associated gene *TSC22D2*^27^ (rs1584586, 3q25.1) with the pathway of purine processing (*PWY-841*, *p* = 3.47×10^-8^). Four suggestive associations for taxa were identified, including signals in 5q35.3 (in the *COL23A1* gene), 8p22 (in the *SGCZ* gene), 14q32.32 (in the *TNFAIP2* gene) and Xp21.1 (in the *DMD* gene).

We sought to replicate our genome-wide significant signals using summary statistics from other independent cohorts in which microbiome data was characterized using either 16S rRNA (MiBioGen study) ^13^ or metagenomic sequencing (LL-DEEP cohort) ^17^. In the MiBioGen study, a meta-analysis of 24 cohorts comprising up to 18,340 individuals, 16S rRNA measurements do not allow for the evaluation of abundance of bacterial species and pathways and the X chromosome was not analyzed. Consequently, only 10 of our 18 SNP‒taxa pairs could be tested in MiBioGen, while none of the pathways were testable. In LL-DEEP, a genome-wide microbiome association study on 952 individuals, we extracted information for the majority of the SNP‒taxa pairs (14/18) and SNP-pathway associations (14/15). Unfortunately, the power to replicate the associations in LL-DEEP was limited due to the small sample size. In both studies, we observed clear replication of the study-wide significant loci, *LCT* and *ABO*. All 7 taxa associated with SNPs near *LCT* were replicated with consistent allelic effect directions (all *p* < 3.7×10^-6^). For the *ABO* locus, we found convincing replication significance for the *Collinsella* genus (*p* < 2×10^-5^ in MiBioGen) and replication at nominal significance for the metabolic pathway *LACTOSECAT-PWY* and *B. bifidum* species (*p* < 0.05 in LL-DEEP) (**Supplementary Table 6** and **Supplementary Table 7**). None of the other SNP‒taxa or pathway pairs were replicated in MiBioGen or LL-DEEP. Interestingly, another independent SNP in the *COL23A1* gene (rs11958296; r^2^ = 0.1 with rs10447306 from our study), shows association in MiBioGen to the abundance of the same taxa: family Rikenellaceae (*p* = 2.4×10^-5^) and genus *Alistipes* (*p* = 9.3×10^-6^).

To explore if the association signals at lower levels of significance are enriched in heritable bacteria, which would indicate if it were possible to detect more genome-wide significant mbQTLs by further increasing the sample size, we investigated the correlation of taxa and pathway heritability estimations gained from family-based analysis with the number of suggestively associated loci for each taxon and pathway. Here we observed a positive significant correlation for both taxonomy (Spearman correlation *r*S = 0.322, *p* = 2.2×10^-6^) and pathway (*r*S = 0.316, *p* = 3.8×10^-6^) heritability and the number of suggestive (*p* < 5×10^-4^) loci identified in our GWAS (**Figure 5**). The correlation remained significant when increasing the mbQTL threshold to *p* < 1×10^-4^ (taxa: *r*S = 0.308, *p* = 6.1×10^-6^; pathways: *r*S = 0.259, *p* = 1.7×10^-4^) and to *p* < 1×10^-5^ (taxa: *r*S = 0.195, *p* = 4.8×10^-3^; pathways: *r*S = 0.148, *p* = 0.034) and when removing the *LCT* and *ABO* loci from analyses (taxa: *r*S = 0.314, *p* = 3.9×10^-6^; pathways: *r*S = 0.311, *p* = 5.7×10^-6^).

**Figure 5.**
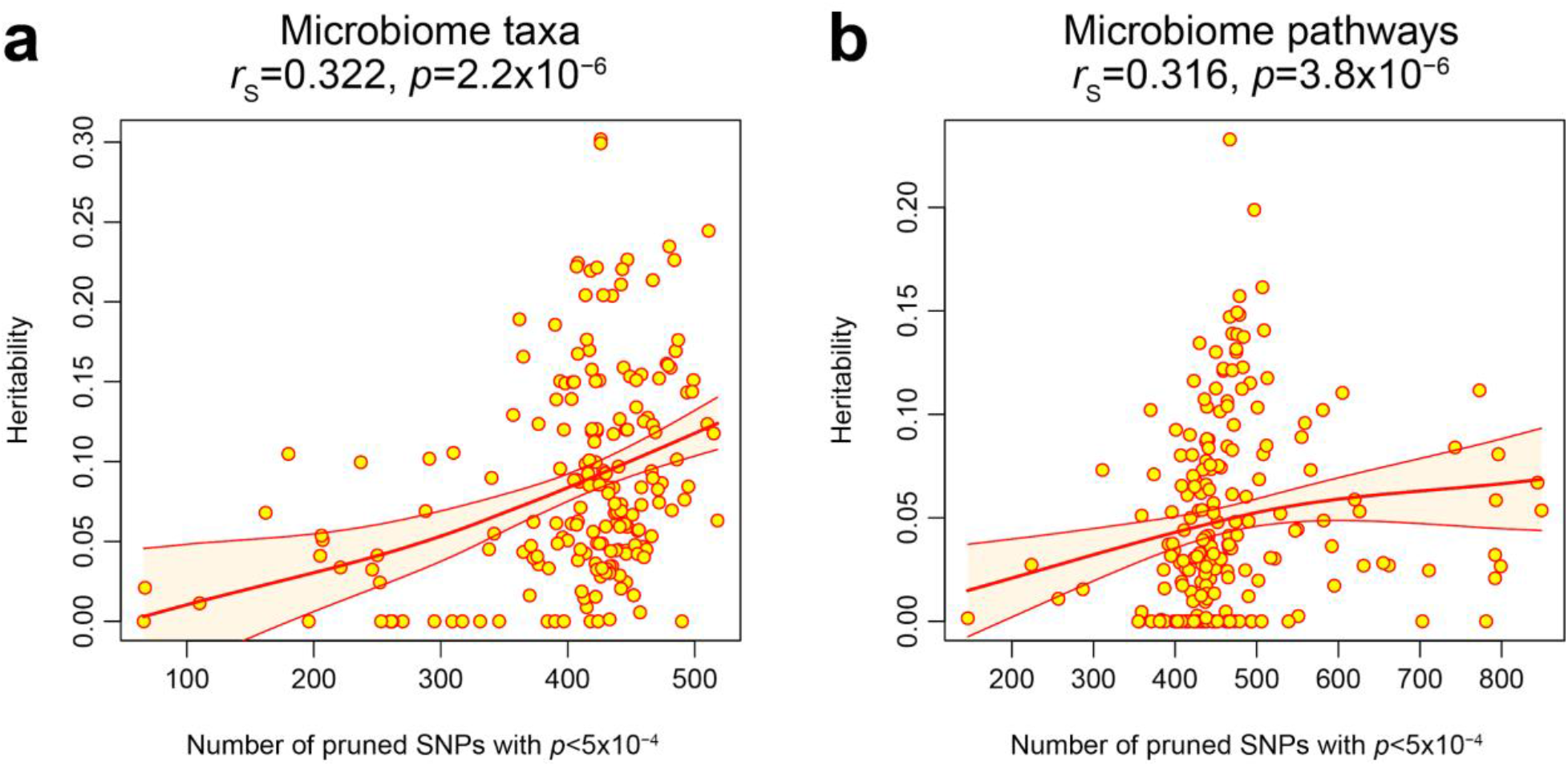
Weighted Spearman correlation between estimated heritability and number of suggestive loci. Each point represents one taxon or pathway. Bold red line represents the locally estimated scatterplot smoothing fit, with orange shaded bands indicating the 95% regression confidence intervals. The number of independent suggestive mbQTLs was obtained using LD pruning (**Methods**). **a)** weighted Spearman correlation of number of suggestive mbQTLs and family-based heritability for microbiome taxa. **b)** weighted Spearman correlation of number of suggestive mbQTLs and family-based heritability for microbiome functional pathways.

We also evaluated if we could replicate any of the genome-wide significant signals outside the *LCT* and *ABO* loci reported in the very recent and similarly sized Finnish population study ^18^. After extracting all associations with *p* < 1×10^-4^ in our dataset, we identified 3 out of 451 genome-wide significant SNPs from the Finnish study using direct or proxy (*r*^2^ > 0.8) information (279 of the SNPs reported in this study were not included in our dataset because their MAF was < 0.05). For one SNP, rs642387, we identified an association with consistent allelic effect for similar taxa: the minor allele at this variant (through proxy SNP rs632222) was associated with a decreased abundance of the *Desulfovibrio* genera (*p* = 8.7×10^-5^) in our study. This corroborates the allelic effects seen in the Finnish study for the abundances of phylum Desulfobacterota A, class Desulfovibrionia, order Desulfovibrionales and family Desulfovibrionaceae. For these taxa, we observed consistent direction in our dataset, although the signals were not significant (*p* = 0.14) (**Supplementary Table 8**).

### Food preference and cardio-metabolic phenotypes link to gut microbiome genetic associations

To investigate the causal relationships between microbiome composition/function and complex traits, and *vice-versa*, we used publicly available summary statistics in conjunction with our mbQTL results to perform Mendelian randomization (MR) analyses (see **Methods**). We focused on 73 phenotypes of autoimmune disease, cardiometabolic disease traits and food preferences (**Supplementary Table 9**) and on 33 microbiome features that were associated with at least one variant at genome-wide significance (*p* < 5×10^-8^). None were significant at FDR < 0.05. At FDR < 0.1 (corresponding to a *p* = 2.7×10^-5^ obtained from the inverse-variance weighted (IVW) MR test), we observed one causal relationship in the direction from microbiome to phenotypes. A genetic predisposition to higher abundance of family Rikenellaceae was causally linked to lower consumption of salt (‘Salt added to food’ ordinal questionnaire phenotype, *IVW p* = 2.4×10^-5^, causal effect = -0.05 in SDs of salt intake for each SD increase in Rikenellaceae abundance) based on three genetic instruments on different chromosomes (**Supplementary Table 10**) (**Supplementary Figure 2a**). This correlation was not significant in the opposite direction (IVW *p* = 0.32), suggesting that Rikenellaceae abundance may have an effect on the increased consumption of salt without being tagged by a shared causal factor. The causal estimate was consistent when using a different MR test (weighted median *p* = 0.00129, causal effect = -0.0554 (SD Salt intake)/(Rikenellaceae abundance SD)). Furthermore, there was no evidence for the causal effect being affected by pleiotropy (Egger intercept *p* = 0.75) or dependent on a single variant (highest *p-value* in a leave-one-out analysis: 8.8×10^-4^) (**Supplementary Figure 1**). At a lower threshold (FDR < 0.2), we observed putative causal effects of two other microbial features on phenotypes: a negative effect of genera *Collinsella* abundance on the level of triglycerides (FDR = 0.10) and a positive effect of the histidine degradation pathway (MetaCyc: “*HISDEG-PWY: L-histidine degradation I*”) on intake of processed meat (FDR = 0.12) (**Supplementary Figure 1**). While all these potential causal effects are intriguing and exhibit robustness to sensitivity analyses, we recognize that the limited availability of genetic variants strongly associated with microbiome features dictates caution in interpretation.

### Power analysis indicates that larger sample sizes are necessary to identify host genetic effects for microbial features with low prevalence

One of the complicating factors in any genome-wide association study of the gut microbiome is the variable detection rate of the pathways and taxa measured, which reduces the effective sample size and consequently statistical power. In our study, the additive effect of associated variants at *LCT* explain 0.8% of the variance for species *B. longum* and *B. adolescentis*, which are present in >80% of the samples, and this allowed us to have ∼70% power to detect the association at *p* < 1×10^-10^ (**Figure 6**). In contrast, we had no power (<5%) to detect the association for species *B. bifidum* and *B. catenulatum* at this level of significance, as they were present in only 14% and 26% of samples, respectively. For *B. bifidum*, despite being a rare species, we were able to detect a study-wide significant association at *ABO* because the effect was substantial (2.5% of the variance explained by SNP rs8176645, power = 83%), but we were underpowered to see the association at this locus with species *C. aerofaciens*, which was present in only 45% of the samples, and for which the additive effect of ABO variants was smaller (0.5% variance explained) (**Figure 6**). In a power analysis, we estimated that our sample size (7,738 individuals) is underpowered to detect genetic effects for taxa that are present in < 80% of samples when considering an effect size comparable or smaller to the effect of *LCT* for *B. adolescentis.* This is striking, considering that 93% of the taxa we identified in this cohort are present in < 80% of the samples (**Methods**). We estimate that to find a similar effect size for at least 20% of gut microbiome composition (approximately 20% of the taxa are present in at least 20% of the samples in our cohort) would require a cohort of ∼50,000 individuals.

**Figure 6.**
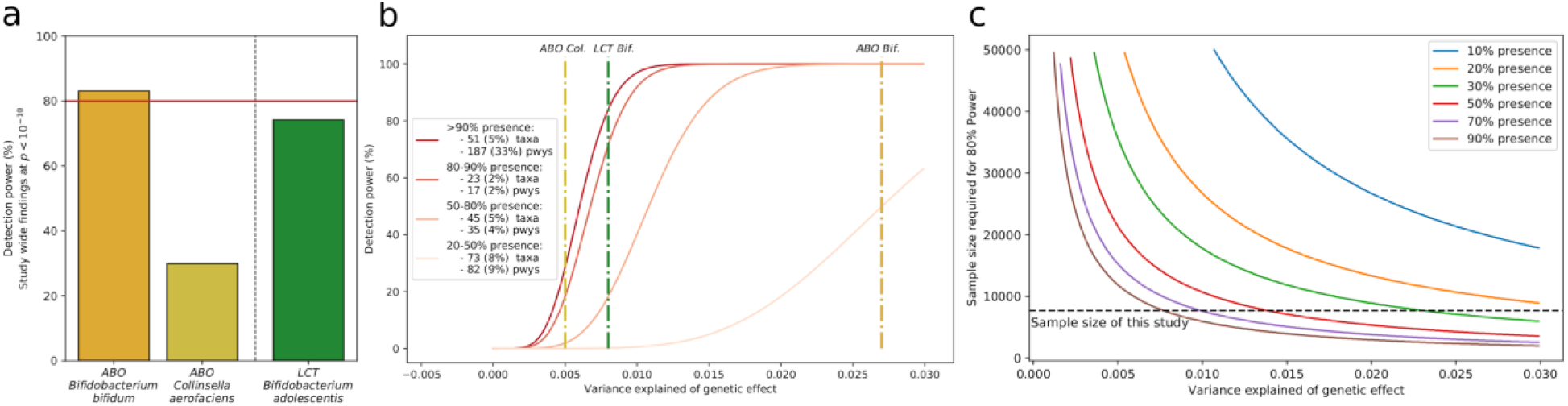
Power analysis taking into account bacterial presence levels. **a)** Statistical power (at alpha = 1×10^-10^) in our dataset for SNPs at the *ABO* and *LCT* loci, considered the associated taxonomies. 80% detection power is shown by the red horizontal line. **b)** Power to detect an effect (y axis, alpha = 1×10^-10^) dependent on the variance explained by a genetic effect (X-axis) in our study. Colored lines distinguish power under certain levels of bacterial or pathway (pwy) presence, which reduces effective sample size. **c**) Sample size needed to identify an association at 80% power level across different bacterial presence levels and alpha = 1×10^-10^. Horizontal line depicts the sample size of the present study.

## Discussion

We carried out the largest genome-wide association study of the gut microbiome composition and function in a single population by analyzing metagenomic sequencing data in 7,738 volunteers from the northern Netherlands. This recapitulated genetic associations at two known loci, *LCT* and *ABO*, and highlighted novel associations with species and bacterial pathways in these loci. Furthermore, we identified associations (*p* < 5×10^-8^) at 18 other loci for 4 taxa and 14 bacterial pathways. None of these associations were affected by major confounders of the gut microbiome such as drug usage, diet and BMI. Finally, we used an MR approach to pinpoint potential causal links between gut microbiome composition, complex traits and food intake habits.

The strongest association with the gut microbiome was found at the *LCT* locus, which remains the most robust genetic association with gut microbiome identified to date. Associations at this locus with *Bifidobacteria* have been consistently reported in studies of different ethnicities, across a range of sample sizes, and in studies using different technologies and protocols for gut microbiome characterization^6, 8, 13, 14, 28^ . We also confirmed that the increase of *Bifidobacterium* was more evident in LI individuals who were consuming milk or milk-derived products^13, 29^. In addition, given that the resolution of metagenomic sequencing allows for species-level characterization of microbiome profiles, we could show that this effect was mainly attributable to species *B. longum, B. adolescentis, B. catenulatum* and *B. bifidum*, an observation that was corroborated by two very recent studies in Finnish and US Hispanic/Latino populations ^18, 30^.

We found another convincing association in the *ABO* gene locus that was responsible for changing abundance in several taxa and pathways. Associations with microbiome composition at the *ABO* locus have been reported previously for populations of different ethnicities and in non-human species. For example, associations with *Bacteroides* and *Faecalibacterium* were reported in a study of five German cohorts that used 16S rRNA sequencing for gut microbiome characterization^28^ and with the microbial pathway of lactose and galactose degradation in a cohort of 3,432 Chinese individuals ^31^. The importance of *ABO* in determining host‒microbiome interaction has also been recently reported in pigs^32^ . A deletion at this locus that inactivates the ABO acetyl-galactosaminyl-transferase has been shown to change the porcine microbiome composition by altering intestinal N-acetyl-galactosamine concentrations and consequently reducing the abundance of *Erysipelotrichaceae* strains, which have the capacity to import and catabolize N-acetyl-galactosamine ^33^. In our analysis, the strongest association was with *B. bifidum*, which has not been reported before, and with the *Collinsella* genus. We did not detect any evidence of interaction with diet at this locus, although this could be due to limitations in available information as the recording of dietary information was done at different times than stool collection time. We did, however, confirm that association at this locus depends on secretor status determined by the *FUT2* gene^28^ and thus on the ability to incorporate antigens into bodily fluids that are released in the gut. Intriguingly, we observed that taxa associated with the *ABO* locus also showed evidence of association at the *LCT* locus, indicating a common action of these two loci in contributing to the growth of these bacteria. The most compelling hypothesis is that the availability of sugars in the gut, via undigested lactose in LI individuals or secretion of glycans in non-O blood type secretors, provides direct energy sources for these bacteria. This is further supported by the observation that LI individuals and non-O blood type secretors were both associated with increased abundance of a bacterial pathway for lactose and galactose degradation. This mechanism would not explain however the opposite pattern of association at ABO for *B. bifidum*.

The strongest mbQTLs we identified reside in genes under selective pressure. The *LCT* gene is a highly differentiated gene among human populations due to positive selection conferred by the lactase persistence phenotype. It has been estimated that strong selection occurred within the past 5,000–10,000 years, consistent with there being an advantage to lactase persistence and the ability to digest milk in the setting of dairy farming ^34^. Variants at this locus have been linked through GWAS to not only food habits and metabolic phenotypes, but also to immune cell populations ^35^. The *ABO* locus is evolutionarily highly differentiated; it has been shown to have experienced balancing selection for the last 3 million years in many primate species ^36^. Several evolutionary sources of selective pressure have been proposed, including via infections by pathogens such as malaria^37^ and cholera^38^. *ABO* variants have also been linked to cardiometabolic traits, white and red blood cells and cytokine levels^39, 40^. Therefore, host‒ microbiome interactions are likely to be shaped by human‒microbe co-evolution and survival, probably through a balance between food availability for gut bacteria and enhanced immune response of the host.

From this perspective, it will be crucial to compare genetic studies of the gut microbiome in diverse populations with different genetic backgrounds in order to understand the complexity of host‒ microbe interactions. This requires community efforts to standardize definition of taxonomies and measurement methodologies in order to facilitate comparison between cohorts. For example, in our attempt to replicate the findings from the Finnish population cohort, only a limited number of taxa could be directly matched or connected through the Genome Taxonomy Database ^41^.

To explore the causality of the relations of the microbiome with complex traits and food preferences, we performed bi-directional MR analysis using 56 dietary traits, 16 diseases and 5 biomarkers. At FDR < 0.2, we observed three causal relationships. Although the limited impact of genetic variants on both microbiome composition and dietary preferences requires caution when interpreting causality estimation by MR^42, 43^, it is intriguing that two observations indicate the causal role of microbial composition on food preferences. We observed that an increase in Rikenellaceae abundance led to decreased consumption of salt and an increase in the bacterial pathway of histidine degradation led to increased intake of processed meat. Although multiple studies have shown that dietary changes, such as variations in salt and meat intake, have a strong effect on microbiome composition^44^, it is intriguing to suggest that genetically determined variations in the microbiome might affect individual food preferences. This would be supported by bacterial genetic variations in salt tolerance and carnitine metabolism genes^45^ and by the established knowledge that the composition of gut microbiome can accurately predict the effect of food items on host metabolism^46^. There was additional evidence from our results to support a role for the microbiome in influencing food preferences. A perfect proxy of rs642387 (rs503397, r^2^ = 0.99 in 1000 Genomes Europeans) that was recently reported to be associated with the family Desulfovibrionaceae and related taxa in a Finnish population, and replicated in our study, had been previously associated to bitter alcoholic beverage consumption in an independent cohort^47^. Another study found an increase in family Desulfovibrionaceae in individuals with high alcohol consumption^48^, and family Desulfovibrionaceae, genus *Desulfovibrio* and other related taxa were also associated with increased consumption of alcohol in our DMP cohort^16^, supporting a pleiotropic effect of this locus on both microbiome and alcohol intake. These studies, together with our findings, suggest that gut microbiota could indirectly influence an individual’s food preferences by mediating the downstream effect of consumption of different products.

Understanding the differences in microbial pathway abundance is crucial for deeply understanding the underlying function of the gut microbiome. Microbiome measurements that target single organisms or taxonomic groups may be inefficient because two strains of one species may differ in their functional capacities. It is therefore essential to evaluate not only the taxonomic but also the functional composition of the microbiome. To our knowledge, ours is the largest single study investigating host genetics effects on the human gut microbiome function using metagenomics sequencing, and thus perform genome-wide association analysis of microbial pathways. Our analyses revealed several loci associated with microbial pathways, although only one passed stringent study-wide significance.

We also observed 18 loci at genome-wide significance and many more at a more lenient threshold. The correlation we observe between the heritability of microbial taxa and pathways and the number of suggestively associated loci indicates that mbQTLs with smaller effects are likely to exist. Those loci remained under the detection limit in the current study, and the sample size would need to be increased by orders of magnitude to discover their role. In fact, gut microbiome composition is characterized by high inter-individual variation. Analysis of core microbiota in different populations indicates that only a few bacteria are present in >95% of studied individuals ^5, 13, 16^, drastically reducing the effective sample sizes for analyses. We performed power calculations taking into account the inter-individual variations in microbiome composition and concluded that a sample size comparable to our study (∼8,000 participants) is only sufficient to identify associations to taxa present in >80% individuals, which comprise only 7% of all identified taxa in the cohort. For bacteria present in >20% of the samples, >50,000 participants would be necessary to identify an effect size similar to that of *LCT* and *ABO*. Much larger cohorts are required to identify genetic contribution to rarer bacteria. We conclude that joint efforts that combine tens of thousands of individuals, combined with harmonized methodology to reduce the technical bias, will be needed to characterize more than a few major loci, as has also been the case for genetic studies of much more heritable quantitative traits such as BMI (heritability ∼40%), height (heritability ∼80%) and others^49–51^.

## Methods

### Cohort description

Lifelines is a multi-disciplinary prospective population-based cohort study with a unique three-generation design that is examining the health and health-related behaviors of 167,729 people living in the north of the Netherlands. Lifelines employs a broad range of investigative procedures to assess the biomedical, socio-demographic, behavioral, physical and psychological factors that contribute to the health and disease of the general population, with a special focus on multi- morbidity and complex genetics ^52, 53^. During the first follow-up visit, a subset of 8,208 Lifelines participants were enrolled in a parallel project, The Dutch Microbiome Project (DMP).The goal of this cohort is to evaluate the impact of different exposures and life-styles on gut microbiota composition^16^.

The Lifelines study was approved by the medical ethical committee from the University Medical Center Groningen (METc number: 2017/152). Additional written consents were signed by the DMP participants or legal representatives for children aged under 18.

### Genome characterization

Genotyping of 38,030 Lifelines participants was carried out using the Infinium Global Screening array® (GSA) MultiEthnic Diseases version, following the manufacturer’s protocols, at the Rotterdam Genotyping Center and the department of Genetics of the University Medical Center Groningen. Here, we used available quality controlled genotyping data imputed with Haplotype Reference Consortium (HRC) panel v1.1^54^, as described elsewhere^55^. For this study, we analyzed European samples, identified through a Principal Component Analysis (PCA) that projected 43,587 genetic markers previously pruned from our genotyping data (sliding window of 1Mb, linkage disequilibrium *r*^2^<0.2, step = 5) onto the 1000 Genomes (all populations) and GoNL cohort as a population reference ^56, 57^. Europeans were identified as samples clustering together with European populations according to the first two principal components (<3 standard deviations from any of the 1000G European samples or GoNL samples), and 35 non-European participants were removed. Quality-controlled genotype information was obtained for 7,738 of the DMP participants for whom quality-controlled microbiome data and BMI were also available. Of these, 58.1% were females and had ages ranging from 8 to 84 (mean of 48.5 years). The mean BMI value was 25.58 (range from 13.10 to 63.70).

### Microbiome characterization

The gut microbiome was characterized from stool samples as described in Gacesa et al.^16^: In brief, stool samples were collected by participants and frozen within 15 minutes after production and then transported on dry ice to the Lifelines facility to be stored at -80°C. Microbial DNA was extracted using the QIAamp Fast DNA Stool Mini Kit (Qiagen, Germany), following the manufacturer’s instructions. Shotgun metagenomic sequencing was carried out using the Illumina HiSeq 2000 platform at Novogene, China. Metagenomic sequencing data was profiled following methods previously used in other cohorts, as described in ^16, 58^. Low quality reads (PHRED quality ≤ 30), adapters and host sequences were removed using KneadData tools v0.5.1. Taxonomic composition was determined with MetaPhlAn2 v2.7.2 ^59^. Characterization of biochemical pathways was performed with HUMAnN2 pipeline v0.11.1^60^, integrated with the UniRef90 v0.1.1 protein database^61^, the ChocoPhlAn pangenome database and the DIAMOND alignment tool v0.8.22^62^. After quality control (samples with eukaryotic or viral abundance ≤ 25% and total read depth ≥ 10 million were retained), we had information on 950 microbial taxa and 559 functional pathways. For this study, we focused only on bacterial taxa and functional pathways with mean relative abundance >0.001% that are present in at least 1000 of the 7,739 participants, which resulted in a list of 207 taxonomies (5 phyla, 10 classes, 13 orders, 26 families, 48 genera and 105 species) and 328 bacterial pathways. Furthermore, we removed redundant pathways by discarding one pathway among pairs that were highly correlated (Spearman *r*^2^ > 0.95), as well as pathways not previously described in bacteria that could thus be coming from sources other than bacteria, resulting in 205 pathways for genetic analyses.

### Diet phenotypes definition

Dietary habits were assessed using a semi-quantitative Food Frequency Questionnaire (FFQ) designed and validated by the division of Human Nutrition of Wageningen University as described before in Siebelink et al and Gacesa et al.^16, 63^. The FFQ data was collected 4 years prior to fecal sampling, and supplementary questionnaires were collected concurrent with fecal sampling, with the stability of long-term dietary habits was assessed as described in Gacesa et al.^16^. We analyzed the dietary factors that were previously found to be associated (False discovery rate (FDR) < 0.05) to the microbiome features in our study that had a genome-wide significant signal in the *ABO* and *LCT* loci. We also analyzed lactose intake for the species *Bifidobacterium longum* and *Bifidobacterium adolescentis*, given their association with the *LCT* region in our study. Participants with an implausible caloric intake (< 800 or > 3934 kcal/day for males, < 500 or > 2906 kcal/day for females)^64^ were not included in these analyses.

### GWAS analysis method

Genome-wide association analysis was performed in 7,738 European samples for 412 features (205 functional pathways and 207 microbial taxa), investigating genetic additive effects using allele dosages for 5,584,686 genetic variants with minor allele frequency (MAF) > 0.05 and information score > 0.4 on the autosomes (chromosomes 1-22) and the X chromosome. We focused on the quantitative dimensions of relative bacterial abundance, treating all zero values as missing data. We used log-transformed taxonomies and regressed these in a linear mixed model using SAIGE v.0.38^19^, with age, sex and the genetic relationship matrix (GRM) among participants as covariates. We used the standard settings of SAIGE, which applies inverse-rank normalization to the traits prior to the association analyses. The genetic relationship matrix was built using a set of 54,565 SNPs selected from the total set of quality-controlled SNPs directly genotyped and filtered for allele frequency and redundancy (MAF ≥ 0.05, r^2^ < 0.2, sliding window = 500Kb). For pathways, we first screened associations with SAIGEgds, an implementation of SAIGE in a computationally efficient framework that allows faster analyses by using best-guess genotypes instead of dosages^65^. We then recalculated genome-wide associations statistics using SAIGE v.0.38 for the 60 pathways that showed at least one genetic variant at *p* < 1x10^-7^.

### Definition of the study-wide significant threshold

To estimate the number of independent phenotypes assessed, we used principal component analysis on the matrix of 412 microbiome features (207 taxonomies and 205 pathways) available for GWAS analysis to decompose variability in independent components (axes). We estimated that 264 components are needed to explain 90% of the microbiome variance. We then defined our study-wide significant *p*-value threshold by correcting the genome-wide significance threshold for this factor (5x10^-8^/264 = 1.89×10^-10^).

### Association using dominant model

To evaluate association using a dominant model on SNP rs182549 at the *LCT* locus, we used best-guess genotypes and converted T/C to T/T. Association analysis was then run for all taxa as well as the “*LACTOSECAT-PWY*” pathway using SAIGEgds, and the same covariates and transformation used for the GWAS analysis.

### Inference of blood groups

We estimated blood groups from genotyped and imputed data following the scheme of Franke et al, 2020^66^. Specifically, we used the absence of the rs8176719 insertion to define blood-type allele O1, the T allele of rs41302905 to define blood-type allele O2 and the T allele of rs8176746 to define the blood-type allele B (instead of rs8176747). Diploid individuals O1O1, O2O2 and O1O2 were considered blood type O. Diploid individuals O1B, O2B and BB were considered blood type B. Absence of the alleles mentioned above was used to define blood-type allele A. To evaluate differences across blood types, we compared the mean relative abundance of microbiome features in individuals with A, B and AB blood type to that in individuals with the O blood type using a two-sided Wilcoxon test. To evaluate the interaction with rs601338 *FUT2* (secretor/non-secretor) locus, we grouped individuals in two groups (non-O blood type and blood type O) to distinguish production or non-production of antigens, and compared pairs using a two-sided Wilcoxon test. All analyses were done using base R v3.6.1 (https://www.R-project.org/).

### Effects of intrinsic and external factors on significantly associated loci

We evaluated the robustness of genome-wide associated signals by incorporating the following potential confounders into our statistical model: medication usage, anthropometric data and stool frequency and consistency data (collection and processing was described in Gacesa et al.^16^). We analyzed the effects of the following drug groups: proton pump inhibitors (ATC A02BC, N = 130), laxatives (osmotic ATC A06AD, N = 44; volume increasing ATC A06AC, N = 77), one group of anti-bacterials (ATC J01, N = 24) and other anti-infectives (ATC J, N = 39). The other group of drugs considered was antibiotic use in the 3 months prior to stool collection (N = 450). For each of these medications, we created dichotomous variables for all participants coded as 0 (non-user) or 1 (user). The list of intrinsic factors included BMI, stool frequency and stool consistency (mean Bristol stool scale). All models also incorporated age and sex as covariates and were run only for the genome-wide‒significant SNP‒trait pairs (**Supplementary Table 1**) using the same software used for GWAS (SAIGE, Zhou et al. ^19^). To evaluate the impact of these covariates on the genetic signals, we used Cochran’s Q heterogeneity test to compare the effect size obtained by the covariate-inclusive model and the basic model (that only includes age, sex and the genetic variant). To evaluate the impact of sex, we ran the SNP‒association analysis in SAIGE separately for males and females using only age as a covariate. For each genetic variant, differences in effect size in males and females were tested using Cochran’s Q heterogeneity test.

### Interaction analyses

We used a three-step procedure to evaluate gene‒diet interactions for all the taxa and pathways associated with SNPs at the *LCT* and *ABO* loci. First, we extracted the variables representing dietary habits that had previously shown significant association with these microbial traits^16^. Next, we added these variables to the basic genetic model (feature ∼ age + sex + genetic variant) to confirm their association at (at least) nominal significance level (*p* < 0.05) while accounting for the associated genetic variant(s). Finally, for the dietary variables showing nominal significance, we evaluated the interaction with the genetic variant(s) by including an interaction term into the association model. For the *LCT* locus, we used SNP rs182549 and considered two groups of genotypes (C/C vs. C/T and T/T) according to the dominant inheritance model. For the *ABO* locus, we used a binary definition of blood type (blood type O and A/B/AB), and also considered the effect of the rs601338 genotype in the *FUT2* gene (defining secretor/ non-secretor individuals).

All microbiome features were inverse-rank normalized before analyses, and age and sex were added as covariates, as in the main GWAS analysis. All models were fit using base R v3.6.1, function *lm()*, statistical tests were as implemented in packages *rstatix* v0.5.0 and *ggpubr* v0.3.0, and package *RNOmni* v0.7.1 was used for the inverse-rank normalization.

### Replication in other cohorts and data sets

We looked for replication of our results using summary statistics from two independent studies: a genome-wide meta-analysis of 16S rRNA data from 24 cohorts and a genome-wide study on metagenomics data in the LL-DEEP cohort, another subset of the Lifelines cohort with data generated 4 years before the DMP and in which 255 participants were also later enrolled in DAG3^13, 17^. In the first study, we could not look at SNPs associated with species or pathways, as these microbiome features cannot be defined using 16S data, or SNPs on the X chromosome, as they were not analyzed in this study. In the second study, we searched for the exact same taxonomy or pathway, but similarly to MiBioGen, X chromosomal variants were not tested and some taxonomies were not defined due to the differences in metagenomics data processing pipelines.

Next to the replication of our findings, we also evaluated whether the genome-wide signals reported in a recent genome-wide study of microbiome taxa from a Finnish cohort were replicable in our data^18^. We searched for all SNPs outside the *LCT* and *ABO* loci or any proxy (r^2^>0.8) in our dataset and selected all SNP‒taxonomy pairs that showed a *p* < 1 x10^-4^ with at least one taxonomy in our cohort (**Supplementary Table 8**). We then looked at these associated taxa in the respective cohorts and compared them visually and with the aid of the Genome Taxonomy Database (https://gtdb.ecogenomic.org/) to determine if they were the same bacterial taxa or taxa from the same taxonomic branch.

### Correlation between heritability estimates and the number of associated loci

To analyze the correlation between family-based heritability and the number of suggestive mbQTLs, we used heritability estimates previously derived from this cohort^16^. We then calculated the number of independent mbQTLs per microbial trait by performing linkage disequilibrium pruning (r^2^ < 0.1 in our data set, window size 1Mb using Plink v1.9^67^) for all SNPs at the three different thresholds: *p* < 5x10^-4^, < 1x10^-4^ and < 5x10^-5^. The association of heritability and the number of mbQTLs was calculated in R v.4.0.3 using a weighted Spearman correlation from ‘wCorr’ v1.9.1 package, with each taxon or pathway treated as a data point. The weights used in calculating the correlation were inversely proportional to the standard errors of heritability estimates. The regression lines in **Figure 5** were fit using the *loess* function (locally estimated scatterplot smoothing, base R package v4.0.3) with span and degree parameters set to 1.

### Mendelian randomization analysis

To evaluate causal relationships between the gut microbiome and other common traits, we performed Mendelian randomization (MR) analyses that combined the summary statistics of the microbiome with publicly available summary statistics on food preferences, autoimmune and cardiovascular diseases and other cardio-metabolic traits. We analyzed the 33 microbiome features (pathways and taxa) with at least one variant passing the *p* < 5x10^-8^ threshold (**Supplementary Table 1**) and combined these with 77 publicly available summary statistic datasets retrieved using the IEU GWAS DATABASE^68^.

We performed a bi-directional MR analysis, first testing if microbiome traits causally affect a phenotype and then testing if phenotypes can causally affect the microbiome traits. For each comparison we intersected the microbiome variants (MAF > 0.05) by rsID, position and alleles with the publicly available summary statistic variants. We then selected instruments using the ‘clump_data()’ function of the TwoSampleMR package (v0.5.5)^69^. The publicly available summary statistics were clumped using a *p-*value threshold of < 5x10^-8^ and otherwise standard settings (r^2^ < 0.001, 10mb window size). Due to the limited statistical significance of the microbiome traits, we performed *p-*value clumping at a less stringent *p* < 5x10^-6^ threshold. If fewer than three variants were clumped, we removed the trait combination from analysis.

The MR analysis was done using the TwoSampleMR v0.5.5 package. We first selected trait combinations that passed the Benjamini-Hochberg FDR threshold of 0.1 (*p* = 2.805x10^-5^) in the inverse-variance weighting test, resulting in one suggestively causal trait combination. We further checked that the significant trait combinations were unlikely to be driven by pleiotropy based on two criteria: i) the Egger regression intercept is nominally significant (indicating the presence of horizontal pleiotropy) and ii) the weighted median results are not nominally significant (indicating that no single variant influences the result)^70, 71^.

### Power analysis

We calculated the variance explained at our loci using the formula described in Teslovich et al. ^72^, which takes into account MAF, effect size, standard error and sample size. We then performed a power analysis based on a linear model of association considering different genetic effect sizes (variance explained) and sample sizes (N) <https://genome.sph.umich.edu/wiki/Power_Calculations:_Quantitative_Traits>. We performed a sample size calculation by doing a grid search in the sample size sequence: [1000, 1050, …, 50000] and keeping the lowest sample size that had power > 80%.

### Data availability

Raw sequencing microbiome data, genotyping data and participant metadata are not publicly available to protect participants’ privacy and research agreements in the informed consent. The data can be accessed by all bona-fide researchers with a scientific proposal by contacting the Lifelines Biobank (instructions at: https://www.lifelines.nl/researcher). The authors declare that all other data supporting the findings of this study are available within the paper and its supplementary information files.

## LifeLines Cohort Study - group authors genetics

Raul Aguirre-Gamboa (1), Patrick Deelen (1), Lude Franke (1), Jan A Kuivenhoven (2), Esteban A Lopera Maya (1), Ilja M Nolte (3), Serena Sanna (1), Harold Snieder (3), Morris A Swertz (1), Judith M Vonk (3), Cisca Wijmenga (1)

(1) Department of Genetics, University of Groningen, University Medical Center Groningen, the Netherlands

(2) Department of Pediatrics, University of Groningen, University Medical Center Groningen, the Netherlands

(3) Department of Epidemiology, University of Groningen, University Medical Center Groningen, the Netherlands

## Acknowledgements

The authors wish to acknowledge the services of the Lifelines Cohort Study, the contributing research centers delivering data to Lifelines and all the study participants. The Lifelines initiative has been made possible by subsidy from the Dutch Ministry of Health, Welfare and Sport; the Dutch Ministry of Economic Affairs; the University Medical Center Groningen (UMCG); the University of Groningen (UG) and the Provinces of the North of the Netherlands (Drenthe, Friesland and Groningen). This project was carried out under Lifelines Project number OV18_0464. We thank Mathieu Plateel and Jody Geelderloos-Arends for their contribution in genotyping the Lifelines samples, Kate McIntyre for editorial support and Patrick Deelen for discussion of results. We also thank the UMCG Genomics Coordination Center, the UG Center for Information Technology and their sponsors BBMRI-NL & TarGet for storage and computational infrastructure.

The generation and management of genotype data for the Lifelines Cohort Study was supported by the UMCG Genetics Lifelines Initiative (UGLI). Metagenomics sequencing of the cohort was funded by the CardioVasculair Onderzoek Nederland (CVON) grant (CVON 2012-03) to MH, JF and AZ. RG and RKW are supported by the collaborative TIMID project (LSHM18057-SGF) financed by the PPP allowance made available by Top Sector Life Sciences & Health to Samenwerkende Gezondheidsfondsen (SGF) to stimulate public-private partnerships and co-financing by health foundations that are part of the SGF. RKW is supported by the Seerave Foundation. AZ is supported by a European Research Council (ERC) Starting Grant 715772, a Netherlands Organization for Scientific Research (NWO) VIDI grant 016.178.056, a CVON grant 2018-27 and the NWO Gravitation grant ExposomeNL 024.004.017. JF is supported by the NWO Gravitation grant The Netherlands Organ-on-Chip Initiative (024.003.001) and CVON grant 2018-27. CW is further supported by an ERC advanced grant (ERC-671274) and an NWO Spinoza award (NWO SPI 92-266). HJMHd is supported by the INTERREG V A (202085) funded project EurHealth-1Health. LC is supported by a joint fellowship from the UMCG and China Scholarship Council (CSC201708320268) and a Foundation De Cock-Hadders grant (20:20-13). EA is supported by Colciencias fellowship ed.783.

## Author Contributions

ELM, AK and SH performed genetic analyses.

AvdG performed Mendelian randomization and power analyses.

AK, SH, LC, AVV and RG processed microbiome data.

SAS, TS, VC, MAYK, LAB and MBG processed samples meta-data.

PBTN and MAS provided resources on the HP computing cluster and assisted with data management.

HJMH, CW, JF, RKW and AZ provided funding resources and designed the DMP.

SZ and SS supervised the study.

ELM, AK, AvdG, SH, SAS, SZ and SS drafted the manuscript.

All authors were involved in data interpretation and provided critical input to the manuscript draft.

## Supplementary Tables Legends

**Supplementary Table 1. Genome-wide significant results**

Table describes independent genome-wide significant results for each taxonomy or pathway. For each variant, we list the rsID, chromosome and genomic position in build 37, reference and alternative allele, alternative allele frequency, number of samples with available information, effect size of the reference allele in standard deviation units (beta) and its standard error, two-sided *p*- value according to the linear mixed model described in the methods, INFO score imputation quality and associated traits. Variants are sorted by *p*-values. Locus ID number identify independent loci, grouping together variants that are in moderate LD (r^2^ > 0.5 in 1000 Genomes Europeans)

**Supplementary Table 2. Association at *LCT* locus under dominant model.**

Table describes association results for the additive and dominant models at SNP rs182549. The effect size (beta) represents the impact of each copy of allele T in the additive model and of at least one copy of allele T in the dominant model, in standard deviation units. Standard errors and two-sided *p*-values are given and were calculated with the linear mixed model as described in **Methods**. For each taxonomy, we also indicate the number of samples with available information and the allele frequency calculated in these samples. Results are shown for all taxonomies that showed a *p* < 1x10^-4^ in the dominant model.

**Supplementary Table 3. Effect of confounding factors at genome-wide significant results.** Table describes the results of sensitivity analyses for the SNP‒trait associations reported in Supplementary Table 1. **Supplementary Table 3a** reports the association results obtained for each SNP at each SNP‒trait association when adding one of the potential confounding factors analyzed as covariate (alternative models). For each alternative model, we report the number of samples analyzed, the effect size in standard deviation units and its standard error, and the Cochran’s Q *p*-value for differences in effect size when compared to the default model. **Supplementary Table 3b** reports association results from the sex-stratified analyses. For each SNP‒trait pair, we report the number of samples (males/females) analyzed, the effect size in standard deviation units and its standard error, and the Cochran’s Q *p*-value for differences in effect size between males and females.

**Supplementary Table 4. Multivariable analysis between diet, microbiome and variants at *LCT* and *ABO* loci**

Table describes the multivariable regression models used to assess the diet effect on microbiome taxa, taking into account age, sex, secretor status, genetic variants at *LCT* and *ABO* loci (**Methods**). The effect size (beta), standard error, t statistics and *p*-value are given for each variable. In addition, the microbial variation explained by all the factors incorporated in models were also provided as adjusted r^2^. The diet factors used (**Methods**; Gacesa et al ^16^) are: average daily Glycemic load (EXP.DIET.Amoungs.GlycLoadTotal), average daily total food intake kcal (EXP.DIET.Amounts.KcalTotal), average daily alcohol intake, energy corrected (g of alcohol per day * [kcal of alcohol / g of alcohol] / total food intake kcal) (EXP.DIET.ECorrected.AlchAll), average daily carbohydrate intake (any carbs), energy corrected (EXP.DIET.ECorrected.CarbsAll), average daily intake of maltose, energy corrected (EXP.DIET.ECorrected.CarbsMaltose), average daily intake of polysaccharides, energy corrected (EXP.DIET.ECorrected.CarbsPoly), average daily intake of lactose, energy corrected (EXP.DIET.ECorrected.CarbsLactose), average daily intake of sucrose, energy corrected (EXP.DIET.ECorrected.CarbsSuchrose), average daily intake of protein (of any type), energy corrected (EXP.DIET.ECorrected.ProtAll), average daily intake of animal protein, energy corrected (EXP.DIET.ECorrected.ProtAnimal), carb to fat ratio = average daily intake of all carbs / average daily intake of all fats (EXP.DIET.Ratios.CarbToFat), carb to protein ratio = average daily intake of all carbs / average daily intake of protein (EXP.DIET.Ratios.CarbToProtein), low carb diet score (EXP.DIET.Scores.LowCarb) with data sorted into 10 deciles based on total carb intake (lowest intake decile = score 10, -1 per decile above it), plant protein diet score (EXP.DIET.Scores.ProtPlantScoredata) with data sorted into 10 deciles based on total carb intake (highest intake decile = 10, -1 per decile below it).

**Supplementary Table 5. Interaction analysis of diet, at *LCT* and *ABO* loci**

Tables describe the interaction analysis between the dietary factors and SNPs at *LCT* or *ABO* loci. For each taxa and pathway, we report the results from the basic models and the results of the models with interaction. A nominal *p*-value < 0.05 of the dietary variable in the basic model was used as a selection criteria for considering this diet factor for interaction analysis (**Methods**). The basic and interaction models for each feature are described in the first column. Sample size, effect size (beta), standard error, t statistics, and *p* value are provided for the variables in each model. An interaction with a nominal *p*-value < 0.05 for the interaction term was considered as a significant interaction effect. The microbial variation explained by all factors incorporated in models is also provided (adjusted r^2^). **Supplementary Table 5a** report results for family Bifidobacteriaceae, **5b** for *B. longum*, **5c** for *B. adolescentis*, **5d** for *B. bifidum*, **5e** for *Collinsella* and **5e** for the LACTOSE-CAT PWY.

**Supplementary Table 6. Replication results in the MiBioGEN cohort**

Table reports the association results listed in Supplementary Table 1, along with the association results obtained previously for the same SNP and taxonomies in the MiBioGen study^13^. For the MiBioGen study, we indicate the number of samples analyzed, *p*-value, effect size in standard deviation units for the alternative allele and its standard error, and the exact taxonomy definition used in MiBioGen study. The last column indicates if the direction of the effect is concordant or discordant with what we observed in our GWAS. SNPs on chromosome X and associations with pathways are not available for MiBioGen.

**Supplementary Table 7. Replication results in the LL-DEEP cohort**

Table reports the association results listed in Supplementary Table 1, along with the association results obtained previously for the same SNP and microbiome feature in the LL-DEEP study^17^. For LL-DEEP, we indicate the number of samples analyzed, *p*-value and effect size in standard deviation units for the alternative allele and its standard error. The last column indicates if the direction of the effect is concordant or discordant with what observed in our GWAS. SNPs on chromosome X and the taxonomies *Collinsella aerofaciens* and *Alistipes spAP11* were not available in LL-DEEP.

**Supplementary Table 8. Replication results of the FINRISK study**

Table reports the association results in our study for SNPs (or their proxy) reported at genome-wide significance in the Finrisk study and located outside the *LCT* and *ABO* loci^18^. For each site, we report the nearby gene, the genomic position of the SNP and the linkage disequilibrium (r^2^) for the proxy SNPs. For each cohort, we report the effect and non-effect allele, the effect size for the effect allele, the standard error, *p*-value and all taxa suggestively associated with the SNP (*p* < 1x10^-4^).

**Supplementary Table 9. Publicly available datasets used in the MR analysis in this study** Descriptions of the publicly summary statistics that were used in the Mendelian randomization analysis of this study. Table contains: the trait ID used to download the data (MRC IEU id), the trait description (trait), year the summary statistics were published (year), (first) author (author), pubmed ID if the study was published (PMID), the unit of the effect size (unit of effect), sample size (sample_size), number of SNPs (nsnps), case-control numbers (ncase if binary & ncontrols if binary), variable type (category) and notes for the phenotype and summary statistics (notes).

**Supplementary Table 10. Mendelian randomization results**

The Mendelian randomization (MR) results of this study. We combined the publicly available summary statistics from **Supplementary table 10** with 33 microbiome features that had at least one significant variant at *p* < 5x10^-8^. We performed MR in both directions (comparison type), comparing the causal effect that an exposure (exposure.id) has on an outcome (outcome.id). We only performed MR analysis if there were >3 instrumental variables available (number_of_instruments). Table shows effect size (beta), standard error (se) and *p*-values (p value) for the inverse-variance weighted (IVW) MR (Weighted_Median, MR_Egger_intercept and MR_Egger_slope). We also list the maximum (least significant) *p*-value in the MR leave-one-out analysis (MR_Leave_One_Out_max_p_value) and the Benjamini-Hochberg false discovery rate *q-*value for the IVW result (q_value). Finally, we list the variants that were used as instruments (instrument_snp_names).

## Supplementary Figures

**Supplementary Figure 1.**
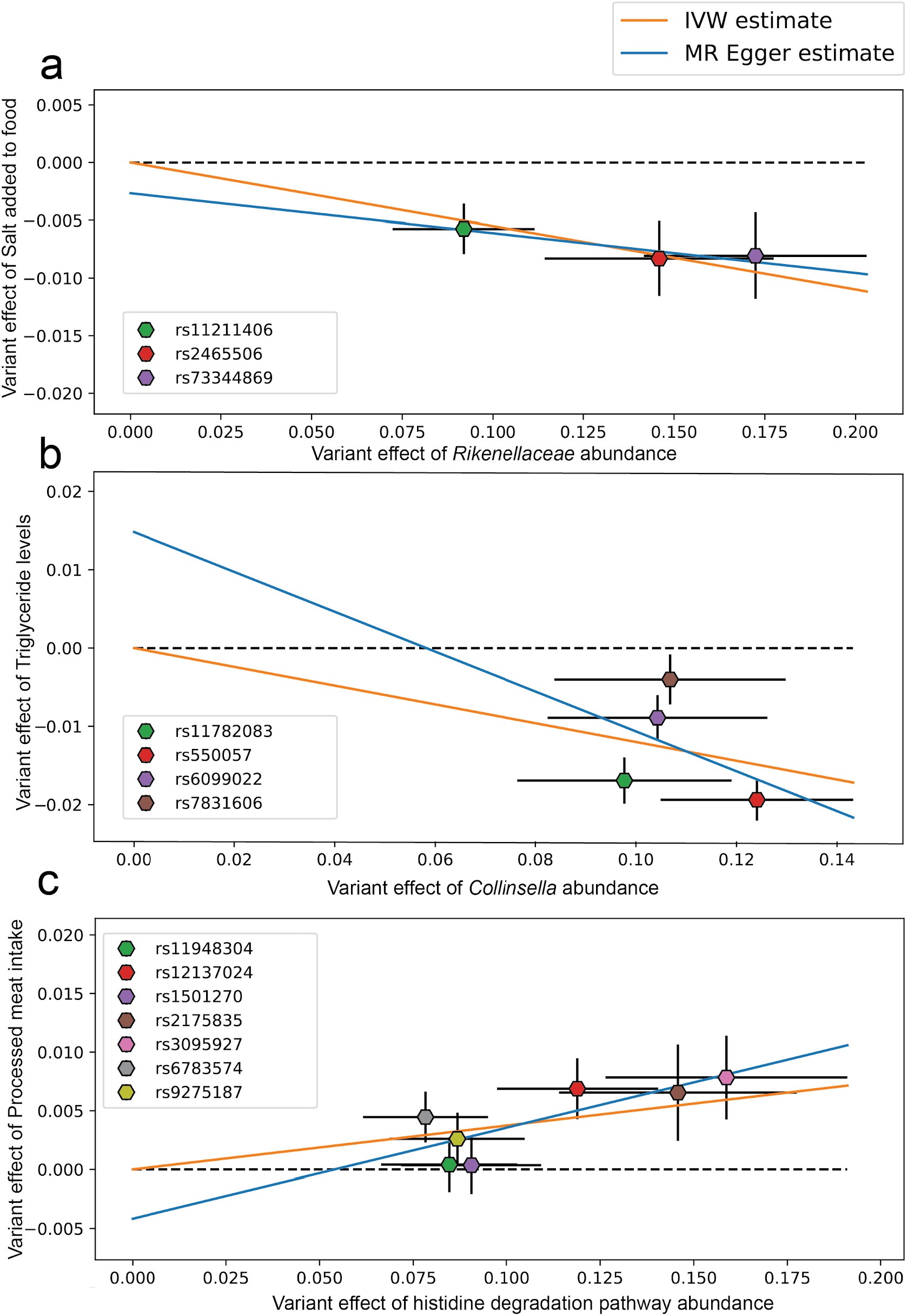
Graphical representation of MR results with a Benjamini-Hochberg FDR q-value < 0.2. **a**) Effect size in standard deviation units of variants associated with Rikenellaceae abundance changes that were used as instrumental variables (X-axis) versus effect size in standard deviation units of the same variants for salt intake (Y-axis). Error bars represent standard errors (SE) of each effect size (beta + SE and beta-SE). **b**) a similar plot to (**a**) but the X-axis is the effect size in standard deviation units for instrumental variants selected for *Collinsella* abundance and on the Y-axis for Triglyceride levels. **c**) effects size in standard deviation units of instrumental variants selected for the abundance of histidine pathway degradation (X-axis) and their effect on the intake of processed meats (Y-axis).

